# *Toxoplasma gondii* from Gabonese forest, Central Africa: first report of an African wild population

**DOI:** 10.1101/2024.05.15.594283

**Authors:** Lokman Galal, Matthieu Fritz, Pierre Becquart, Karine Passebosc-Faure, Nicolas Plault, Larson Boundenga, Illich Manfred Mombo, Linda Bohou Kombila, Telstar Ndong Mebaley, Léadisaelle Hosanna Lenguiyah, Barthélémy Ngoubangoye, Nadine N’Dilimabaka, Eric M. Leroy, Gael Darren Maganga, Aurélien Mercier

**Affiliations:** Inserm U1094, IRD UMR270, Univ. Limoges, CHU Limoges, EpiMaCT - Epidemiology of chronic diseases in tropical zone, Institute of Epidemiology and Tropical Neurology, OmegaHealth, Limoges, France; Maladies Infectieuses et Vecteurs, Ecologie, Génétique, Evolution et Contrôle (MIVEGEC), University of Montpellier, IRD, CNRS, 34394 Montpellier, France; Centre National de Référence (CNR) Toxoplasmose/Toxoplasma Biological Center (BRC), Centre Hospitalier-Universitaire Dupuytren, Limoges, France; Centre Interdisciplinaire de Recherches Médicales de Franceville (CIRMF), Gabon; Department of Anthropology, University of Durham, Durham, United Kingdom; Faculté des Sciences et Techniques, Université Marien Ngouabi, Brazzaville, République du Congo; CNRS, Laboratoire de Biométrie et Biologie Evolutive UMR5558, Université de Lyon 1, 69622 Villeurbanne, France; Département de Biologie, Faculté des Sciences, Université des Sciences et Techniques de Masuku (USTM), Franceville, Gabon; INSAB, Université des Sciences et Techniques de Masuku (USTM), Franceville, Gabon

## Abstract

The protozoan *Toxoplasma gondii* is a ubiquitous and highly prevalent parasite that can theoretically infect all warm-blooded vertebrates. In humans, toxoplasmosis causes infections in both immunodeficient and immunocompetent patients, congenital toxoplasmosis, and ocular lesions. These manifestations have different degrees of severity. Clinical severity is determined by multiple factors, including the genotype of the *T. gondii* strain involved in the infection. *T. gondii* exhibits remarkable genetic diversity, which varies according to geography and ecotype (domestic or wild). Previous studies have demonstrated that wild strains of *T. gondii* are of particular epidemiological interest, as they have been associated with more severe forms of toxoplasmosis in different regions of the world. However, no data on wild strains of *T. gondii* are available from Africa. In this study, we describe for the first time a wild *T. gondii* population from Africa. Wild animals from the forest environment of Gabon, Central Africa, were screened for chronic infection with *T. gondii* using quantitative PCR. The infecting *T. gondii* strains were genotyped whenever possible by the analysis of 15 microsatellite markers and by whole-genome sequencing. A new genotype was identified and was found to be highly divergent from previously described *T. gondii* populations worldwide, including those from the domestic environment in Gabon. Whole genome-based analyses indicated that this strain was genetically closer to a wild Pan-American population than to domestic African populations. This discovery marks the first description of a wild *T. gondii* population in Africa. The role of wild *T. gondii* strains in the incidence of severe toxoplasmosis in Africa remains unclear and requires further investigation.

**Author Summary:** The emergence of new pathogens from wildlife is today a well-recognized health threat. Studying these infectious agents has proven to be challenging due to the difficulty in accessing to samples from wild animals. In the present study, we took advantage of a recent survey on the viral carriage of wild animals from Gabon, Central Africa, to screen animal samples for the presence of the zoonotic protozoan *Toxoplasma gondii*, a ubiquitous and highly prevalent parasite that can theoretically infect all warm-blooded vertebrates, including humans. This parasite is the etiological agent of toxoplasmosis, a disease causing a substantial public health burden worldwide through different clinical manifestations and varying degrees of severity. A novel genotype was identified and found to be highly divergent from previously described *T. gondii* populations worldwide, including those from the domestic environment in Gabon. This discovery marks the first description of a wild *T. gondii* population in Africa. It has been shown that wild strains of *T. gondii* are of significant epidemiological relevance, as they have been associated with more severe forms of toxoplasmosis in different regions of the world. The implications of wild *T. gondii* strains in the incidence of severe toxoplasmosis in Africa remain unclear and merit further investigation.

## Introduction

*Toxoplasma gondii* is a ubiquitous zoonotic protozoan infecting all warm-blooded species, including humans. All these species can act as intermediate hosts for *T. gondii* by developing persistent tissue-cysts after feeding from tissues of another infected intermediate host or following the ingestion of sporulated oocysts found in the environment [1]. These oocysts are excreted in the environment through the feces of members of the Felidae family, the only definitive hosts of this parasite, following their feeding on infected prey [2]. The oocysts sporulate within few days following their excretion to become infective.

*T. gondii* is estimated to infect around 30% of the human population and is the etiological agent of toxoplasmosis, a disease causing a substantial public health burden worldwide [3]. Historically, infection with *T. gondii* has been long considered essentially asymptomatic or benign, except for certain risk groups, such as the developing fetus in the case of congenital infection, and immunocompromised patients for whom toxoplasmosis can have severe health consequences either during primo-infection or reactivation [4]. However, clinical toxoplasmosis has also been reported in immunocompetent individuals, mainly in the form of ocular lesions and multivisceral involvement [5–7]. The prevalence of clinical toxoplasmosis, its clinical forms and their severity substantially vary worldwide [8]. *T. gondii* strains diversity, which exhibit contrasting patterns across geographic regions and ecotypes, appears to explain, at least in part, this clinical variability [9].

In Africa, the few available reports of clinical toxoplasmosis in immunocompetent individuals suggest that certain *T. gondii* strains on this continent are more pathogenic than European strains [10–14]. However, the association between genotype and disease severity is still unclear due to the scarcity of these reports. Furthermore, nearly all *T. gondii* isolates have been collected on human and domestic animals, and no sylvatic cycle of *T. gondii* has been described on this continent. Wild strains of *T. gondii* have been demonstrated to hold significant epidemiological importance, as they have been associated with more severe forms of toxoplasmosis in other regions of the world [15–17].

In the present study, we took advantage of a recent survey on the viral carriage of wild animals from northeast Gabon, central Africa [18]. A total of 148 animals belonging to at least seven species were tested for chronic *T. gondii* infection using a quantitative PCR assay to detect *T. gondii* DNA in different organ samples. The infecting *T. gondii* strains were genotyped whenever possible by the analyzing 15 microsatellite (MS) markers and by Illumina whole genome sequencing (WGS). The objective of this study was to provide the first insights into the *Toxoplasma gondii* strains circulating in African wildlife and to provide evidence for the existence of a sylvatic cycle of *T. gondii* in Africa.

## Material and Methods

### Ethics statement

To carry out the sampling campaign, we obtained authorization to capture and collect animals from the Ministry of Water and Forests, in charge of the environment and sustainable development (Authorization No. 0247 MEFCEDD/SG/DGFAP).

Organ samples (brain, heart, lung and kidney) were previously collected from wild animals hunt in the surrounding forest of 11 villages in the department of Zadié, province of Ogooué-Ivindo, in northeast Gabon in 2019 [18]. Here, these organs were analyzed in order to detect chronic *T. gondii* infection in these animals. For this purpose, around 30 mg of each organ was collected when available and transferred to Lysing Matrix E tubes (MP Biomedicals) at -80 °C until processing. Samples were mechanically disrupted using a TissueLyser II (Qiagen, Courtaboeuf, France) for 30s at 30 Hz. Then cooling of samples was performed in dry ice for 45s, before carrying out second round of mechanical disruption under the same conditions. Tubes were then centrifugated at 200 × g for 5 min and 350 µl of lysate was collected from each tube for DNA extraction using the QIAamp DNA Mini Kit (Qiagen, Courtaboeuf, France).

The extracted DNA samples were tested by a quantitative polymerase chain reaction (qPCR) assay as described by Ajzenberg et al. [19] on a thermocycler Rotor-Gene 6000 (Corbett Life Science, Sydney, Australia), targeting the 529bp DNA fragment (REP529, GenBank accession no. AF146527 [20]). In brief, each PCR contained 5 μL of extracted DNA from organs, mixed with 15 μL of a PCR mix with 1X LightCycler FastStart DNA Master Hybridisation Probes kit (Roche diagnostics, Mannheim, Germany), 0.5 U of UDG (Roche Diagnostics, Mannheim, Germany), 5 mmol/L of MgCl2, 0.5 μmol/L of each primer, 0.1 μmol/L of TaqMan probe (Eurofins, Ebersberg, Germany) which is labeled with a fluorescent dye (6-carboxyfluorescein, 6-FAM) at 5’ end and a dark quencher (Black Hole Quencher, BHQ1) at the 3’ end. The cycling protocol was as follows: initial decontamination by UDG at 50ºC for 2 min and denaturation at 95ºC for 10 min, followed by 50 cycles at 95ºC for 20 s and 60ºC for 40 s. Each sample was run in duplicate and the results obtained were expressed in cycle threshold (Ct) values.

*T. gondii* strains were genotyped using 15 MS markers distributed across 11 of the 13 chromosomes composing *T. gondii* genome in a single multiplex PCR-assay as described previously [21], following the guidelines established by Joeres et al. [22]. Those 15 loci included a combination of eight “typing” markers with low polymorphism (TUB2, W35, TgM-A, B17, B18, M33, IV.1 and XI.1) that show little or no variation within lineages and seven “fingerprinting” markers (M48, M102, N83, N82, AA, N61, N60) exhibiting high polymorphism and significant variation within lineages. PCR products were sized using capillary electrophoresis on ABI PRISM 3130xl (Applied Biosystems, Foster City, CA) and the GenScan 500 ROX dye size standard (Applied Biosystems). Results were analysed using GeneMapper 5.0 software packages (Applied Biosystems). New multilocus genotypes (MLGs) were compared to those from a global dataset (S1 Table) of previously published MLGs (n = 1068) by generating a neighbor-joining dendrogram using the BRUVO.BOOT function (based on Bruvo’s genetic distance) with 1,000 bootstrap replications, as implemented in the “Poppr” package [24] in R version 3.4.0. In addition, the factorial correspondence analysis (AFC) technique available in GENETIX version 4.05 (Belkhir et al. 2003) was used to visualize the genetic distance between MLGs in a multidimensional space (3D).

WGS was performed for DNA samples displaying high *T. gondii* DNA concentration. These DNA samples were subjected to high-throughput sequencing (HTS) on the Illumina NovaSeq 6000 platform (Novogene, United Kingdom). FastQC was applied to analyze reads quality and adapters were trimmed with Trimmomatic v0.39 to truncate low quality reads, filtering for a minimum read length of 36 (parameters: SLIDINGWINDOW: 4:20; MINLEN: 36; TruSeq3-PE-2.fa:2:30:10).

Trimmed reads were mapped using Smalt v7.6 to the newly available genome assembly ASM1945558v1 (GenBank accession code: GCA_019455585.1), obtained using Oxford Nanopore long read sequencing of the *T. gondii* ME49 strain (type II) [23]. Smalt options for exhaustive searching for optimal alignments (-x) and random mapping of multiple hit reads were used. Reads were mapped when mapping base identity was greater than or equal to 80% to the reference genome (-y 0.8), to eliminate spuriously mapped reads mainly from host DNA. Mapped reads were sorted with Samtools 1.11, and duplicate reads were marked with Picard ‘MarkDuplicates’ 2.25. In order to overcome the issue of low genomic coverage associated to uncultured samples — due to the overwhelming proportion of host DNA —, a different approach was adopted to compare the new strain from this study to global *T. gondii* diversity. Trimmed reads were mapped separately to 16 available reference genomes representing 15 of the 16 *T. gondii* haplogroups (hg) previously described worldwide (the 16^th^ haplogroup was not available) [24]. Stringent mapping configurations were applied to map only reads that show a high degree of identity with these reference genomes. Using this approach, we expected that the number of reads mapped to each reference genome would be an indication of the genetic proximity between the new strain from this study and the *T. gondii* haplogroup to which the reference genome belongs.

## Results

The 148 animals included in this study were of at least seven distinct wild mammal species. However, the number could have been underestimated given that only the genus could be determined for 43 individuals. All the four organs considered in this study (brain, heart, lung and kidney) were available —and therefore were tested using qPCR— in 80 animals while between one and three organs were available and were tested in the remaining 68 animals. *T. gondii* DNA was detected in 15 animals belonging to at least four distinct species (*Philantomba monticola, Cephalophus callipygus, Cephalophus dorsalis, Cephalophus* sp. and *Atherurus* sp.) and from six different villages. In the 15 PCR-positive animals, one to three organ types were PCR-positive, while none was found to be PCR-positive for the four organs. Despite this, 11 of the 15 PCR-positive animals were tested for the four organ samples. For each PCR-positive animal, the organ sample showing the lower C_t_ value was selected for MS genotyping. MS markers could be amplified in two samples: (1) Gabon-87_2019-Cephalophus-sp, a heart sample from a *Cephalophus* sp. from Mekouma (12/15) and (2) Gabon-21_2019-*Cephalophus-callipygus*, a heart sample from a *Cephalophus callipygus* from Zoula (1/15). The two samples displayed a novel allele (228 at M102), not previously observed in any of the previously published MLGs, suggesting that they may belong to the same population. Gabon-87_2019-Cephalophus-sp displayed another novel allele (154 at B18). Notably, it shared four of its five amplified typing markers with TgSpEt19, a strain from a sheep in Ethiopia (Table 1).

**Table 1.**
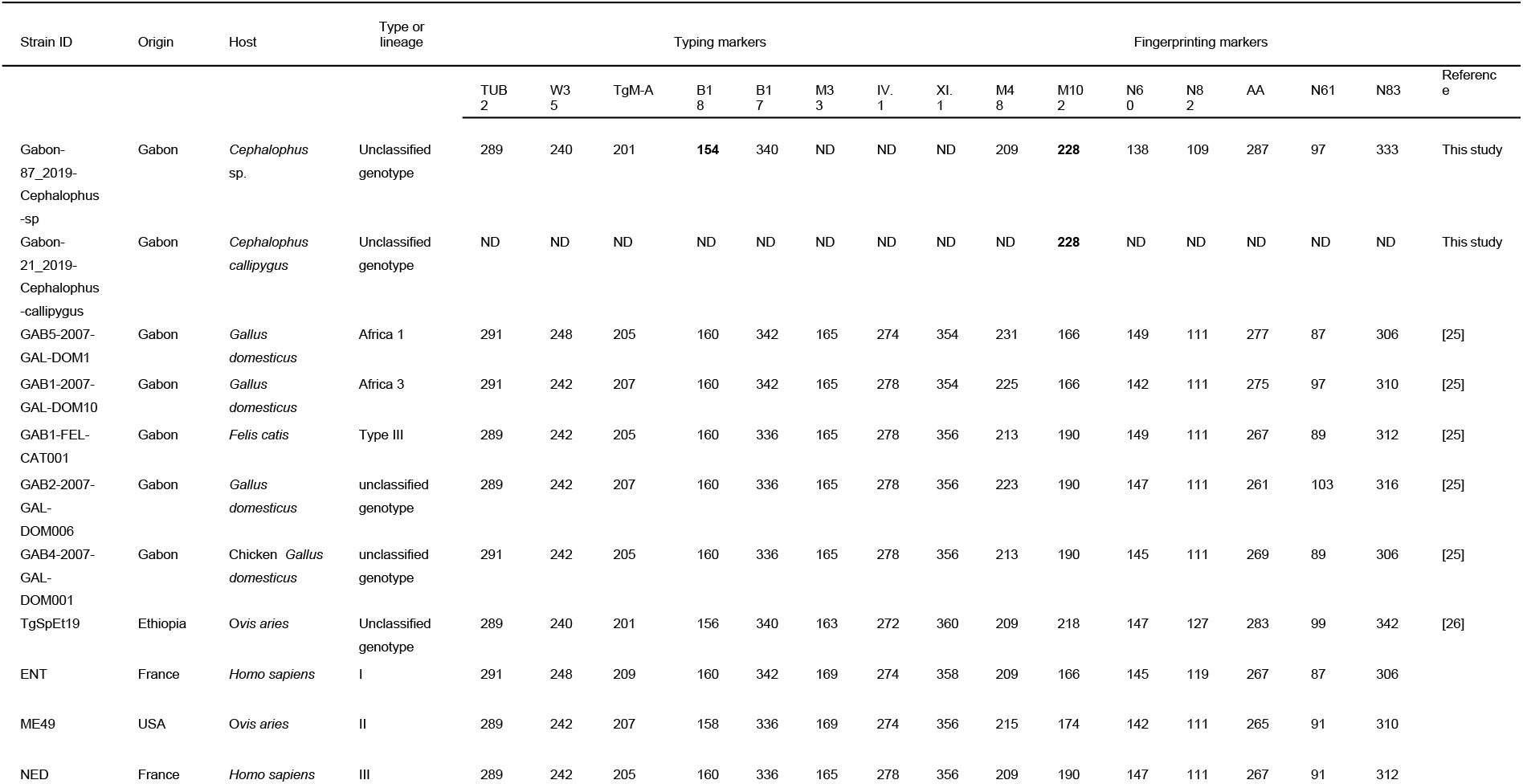
*Toxoplasma gondii* microsatellite (MS) analysis of organ samples from Gabonese wild animals and a comparison with other Gabonese, African and global isolates. New MS alleles are indicated in bold letters.

The NJ dendrogram (Fig.1) and the AFC (Fig.2) confirmed that Gabon-87_2019-*Cephalophus*-sp was genetically related to TgSpEt19, and exhibited divergent profile from other previously described MLGs worldwide, including those from the domestic environment in Gabon.

**Fig. 1.**
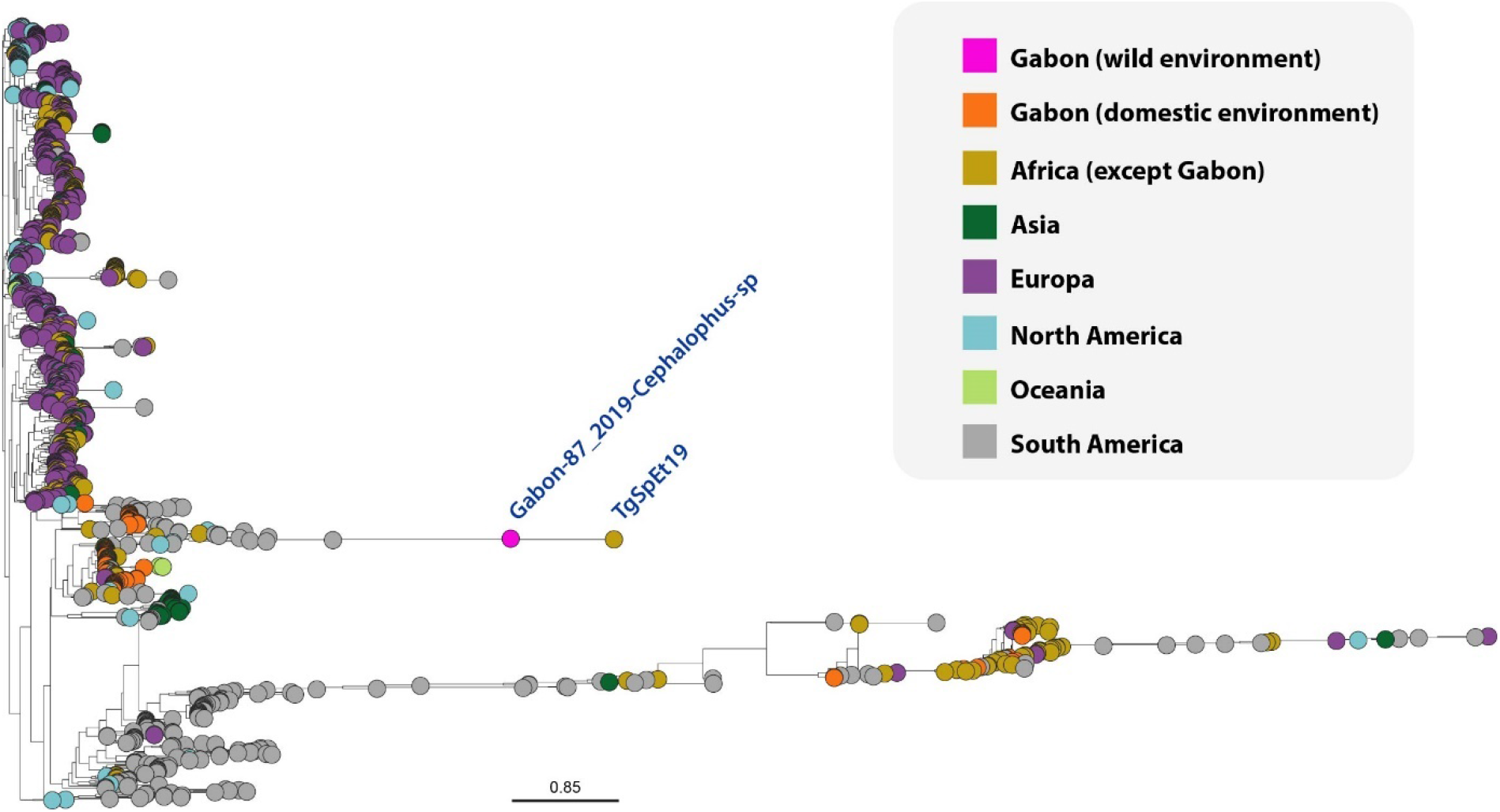
Neighbor-joining tree showing the relationships between Gabon-87_2019-Cephalophus-sp and other global multilocus genotypes (MLGs) (n = 1068) from previous studies.

**Fig. 2.**
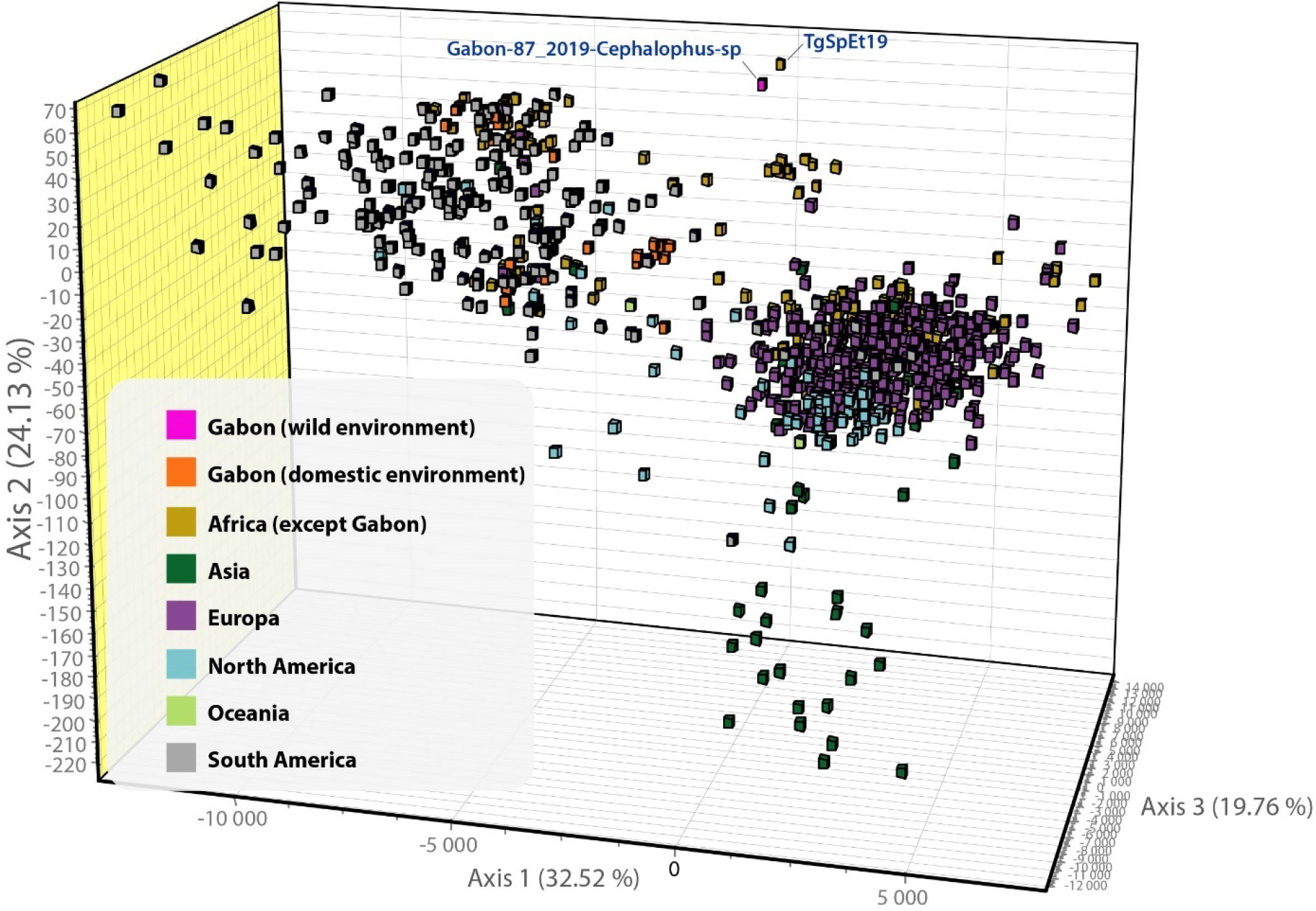
Factorial correspondence analysis (AFC) technique including Gabon-87_2019-Cephalophus-sp and other global multilocus genotypes (MLGs) (n = 1068) from previous studies.

Among the 15 PCR-positive animals, only Gabon-87_2019-Cephalophus-sp (heart sample) was selected for Illumina WGS. Following quality control, 1,371,024,686 paired-end reads (150 nt × 2) were obtained. Only 282,006 (0.02% of trimmed reads) were mapped to the Oxford Nanopore genome assembly ASM1945558v1. The average depth of genome coverage was 3.5, which was insufficient for further analyses. Illumina reads were then mapped to each of the 16 reference genomes representing 15 of the 16 global haplogroups. The reference genome to which the higher number of reads were mapped was COUG (haplogroup 11) (Fig.3).

**Fig. 3.**
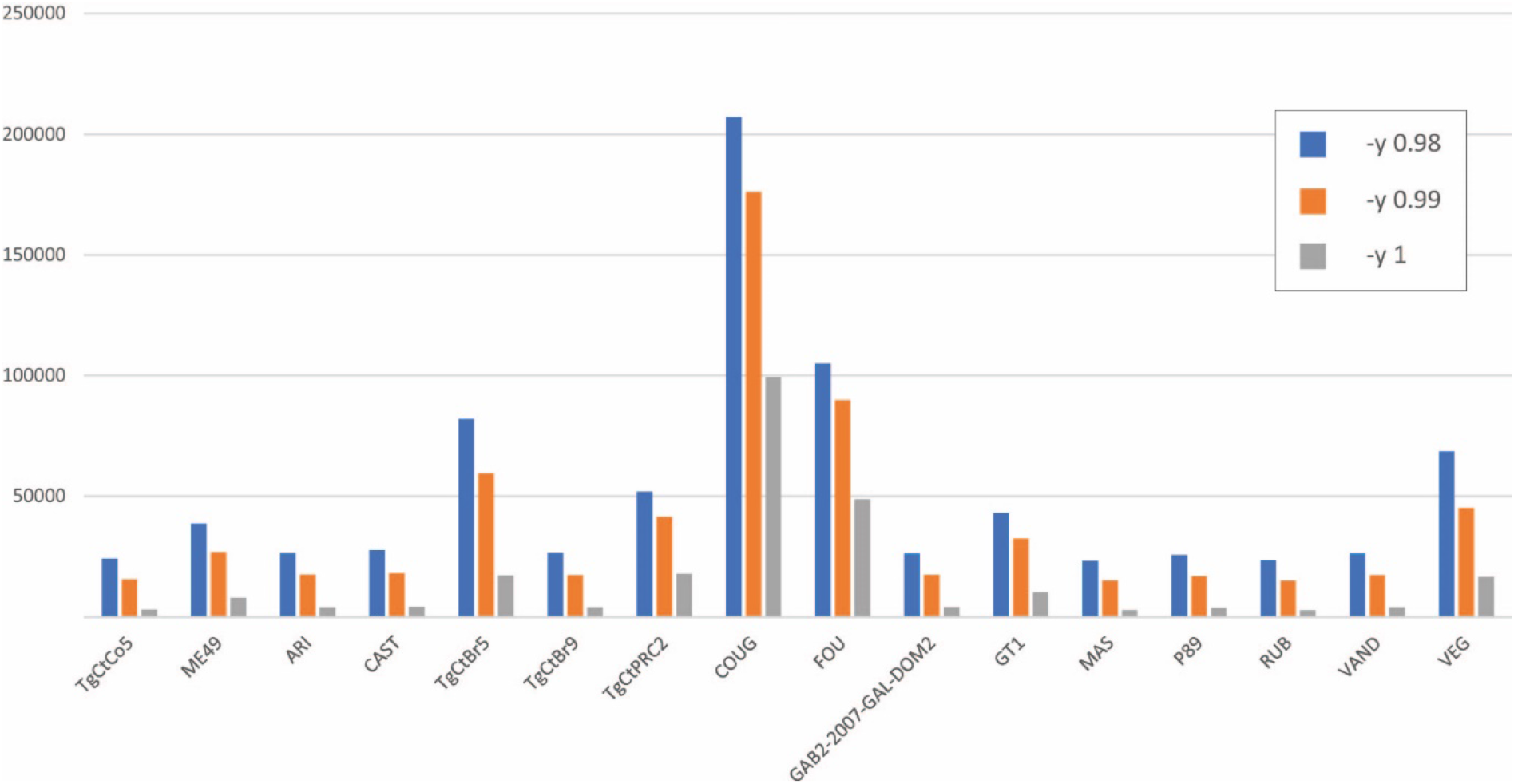
Numbers of Illumina reads from Gabon-87_2019-Cephalophus-sp that mapped to each of the 16 reference genomes. Smalt was executed three times on each of the 16 reference genomes, with a different mapping base identity setting each time (-y 0.98, -y 0.99, -y 1.0). These references genomes are GT1 (type I; hg1; GCA_000149715.2), VEG (type III; hg3; GCA_000150015.2), ME49 (type II; hg2; GCA_000006565.2), MAS (atypical; hg4; GCA_000224865.2), RUB (Amazonian; hg5; GCA_000224805.2), CAST (atypical; hg7; GCA_000256705.1), TgCtBr5 (atypical; hg8; GCA_000259835.1), P89 (atypical; hg9; GCA_000224885.2), VAND (Amazonian; hg10; GCA_000224845.2), COUG (Pan-American; hg11; GCA_000338675.1), ARI (type 12; hg12; GCA_000250965.1), TgCtPRC2 (Chinese 1; hg13; GCA_000256725.1), GAB2-2007-GAL-DOM2 (Africa 4; hg14; GCA_000325525.2), TgCtCo5 (atypical; hg15; GCA_000278365.1), FOU (Africa 1; hg6; GCA_000224905.2) and TgCtBr9 (atypical; hg6; GCA_000224825.1).

## Discussion

This is the first study exploring *T. gondii* circulation in the wild environment of a tropical African country. This study reports for the first time the presence of *T. gondii* in the tissues of species such as duikers (*Philantomba* sp. and *Cephalophus* sp.) and brush-tailed porcupines (*Atherurus* sp.). This study is also the first description of a wild *T. gondii* population in Africa.

To date, the presence of wild *T. gondii* populations could only be confirmed in North America and in French Guiana, in South America [27–30]. These regions are characterised by the persistence of wild felid populations of relatively large sizes [31–33], which are probably the main driver of the maintenance of *T. gondii* sylvatic cycles. In other countries where representative sampling of humans, domestic and wild animals has been carried out, such as China and France, *T. gondii* clonal lineages isolated from all these host categories essentially belonged to the same domestic lineages (mainly Chinese 1 and type I in China and type II in France) [34–37]. In China and France, wild felid populations have undergone a significant decline due to the destruction of their habitats [38,39]. This situation could have resulted in the disappearance of sylvatic cycles of *T. gondii* in these countries, potentially explaining the lack of reports on wild *T. gondii* populations there. Furthermore, a recent study also indicates a lack of ecotype compartmentalization in *T. gondii* populations from Brazil [40], although the numerous gaps in sampling in this country—especially from wildlife—make it challenging to draw definitive conclusions at this stage. It is noteworthy that natural habitats in Brazil are subject to significant degradation in comparison to those neighboring French Guiana [41], which may explain the absence of ecotype compartmentalization. In Gabon, the equatorial forest is one of the few well-preserved ecosystems of its kind in Africa [42] and is a refuge for wild feline species such as the leopard (*Panthera pardus*) and the African golden cat (*Caracal aurata*). This situation may be conducive to the persistence of wild *T. gondii* populations in this ecosystem.

The various patterns of ecotype compartmentalization observed in *T. gondii* populations according to geographical areas have presented a challenge in the assignment of a *T. gondii* strain to an ecotype. In Africa, sporadic reports of *T. gondii* strains from wild animals are available [43,44], but these animals were infected with *T. gondii* strains commonly found in the domestic environment, as it is the case in France and China. The most commonly accepted definition of a wild strain is that it is restricted to the wild environment and genetically distinct from strains found in the domestic environment within the same geographical region [28,30,40]. This is the first time a *T. gondii* population from Africa fits this definition, as it isolated from wildlife and was found to be highly divergent from *T. gondii* populations previously described in domestic animals in Gabon [22]. A recent study on *T. gondii* genomics identified a ∼100-kb genomic region on chromosome 1a that has proven to be a robust marker for distinguishing *T. gondii* strains from different ecotypes (domestic or wild). This study showed that, on a global scale, *T. gondii* strains from the domestic environment display a unique haplotype in this genomic region on chromosome 1a, which is considered a probable specific adaptation to domestic cats. Conversely, wild *T. gondii* strains exhibit a high diversity of haplotypes at this genomic locus [30]. In this instance, the haplotype carried by the wild Gabonese *T. gondii* population identified in this study could not be sequenced due to the low proportion of *T. gondii* DNA in tissue samples and the low coverage of the sequenced genome.

The MS-based analyses revealed a marked genetic proximity between the wild Gabonese *T. gondii* strain Gabon-87_2019-Cephalophus-sp and a unique strain isolated from a sheep in Ethiopia [26]. The flock this sheep belonged to had a grazing area commonly frequented by several species of wild felids [45], which could have been the source of contamination of this sheep. A similar pattern is observed in North America, where grazing domestic animals are substantially more exposed to wild *T. gondii* strains than farm-bound animals [29]. This genetic proximity between Ethiopian and Gabonese strains reveals that certain branches of the *T. gondii* evolutionary tree remain obscure. The global proliferation of domestic cats has favored the spread of few cat-adapted clonal lineages [30], which have probably overwhelmed ancient *T. gondii* populations in many regions of the globe. The massive collapse of most wild felid populations has likely restricted wild *T. gondii* strains to a few relatively well-preserved ecosystems where wild felid populations of considerable sizes are still maintained as observed in French Guiana and in North America [46,32]. This situation has probably facilitated the sampling of wild *T. gondii* strains in these two latter regions whereas wild *T. gondii* strains transmitted by African, European and Asian wild felids remain almost unknown [47]. The inclusion of wild *T. gondii* strains from these regions in phylogenetic analyses could challenge the current paradigm of a South American origin for current *T. gondii* populations [48]. This is particularly relevant given the probable origin of the Felidae family in Asia [49]. In accordance with this, a whole genome-based comparison of Gabon-87_2019-*Cephalophus*-sp with the reference genomes representing the major *T. gondii* haplogroups worldwide suggested that Gabon-87_2019-*Cephalophus*-sp was more genetically related to COUG (haplogroup 11) than to the major domestic *T. gondii* lineages found in Gabon and Africa (type II, type II, Africa 1 and Africa 3). COUG strain has been isolated from a cougar during the investigation of a large community outbreak of waterborne toxoplasmosis in humans in Canada, although the involvement of this strain in human cases could not be demonstrated [50]. COUG strain belongs to a wild *T. gondii* population designated as Pan-American, which displays a divergent haplotype on the ∼100-kb genomic region on chromosome 1a associated to cat adaptation [30]. In addition to Canada, *T. gondii* strains of the same population have been isolated from wild animals in the United States [51], Mexico [52] and French Guiana [28]. This relative genetic proximity between Gabon-87_2019-*Cephalophus*-sp and COUG could reflect an ancient divergence between wild populations of *T. gondii*, probably associated with the history of wild felids global dissemination.

In South and North America, it has been previously shown that *T. gondii* strains from wildlife are often associated with cases and outbreaks of severe ocular and/or systemic disease and unusual presentations of toxoplasmosis in immunocompetent patients [53–56]. These clinical forms have been increasingly diagnosed in the last two decades and are considered to be still underdiagnosed on these two continents. The present study provides evidence for the existence of a sylvatic cycle of *T. gondii* in Africa. The involvement of African wild *T. gondii* strains in the incidence of severe toxoplasmosis among immunocompetent individuals [10–12,14] requires further investigation.

## Supporting information

S1 Table. Global dataset of previously published Multilocus genotypes (MLGs) (n = 1068) obtained from the analysis of 15 microsatellite markers and used for comparison with new MLGs from this study.

## Acknowledgments

We thank Philippe Engandja, CIRMF, Gabon, for their technical assistance during this work. The authors thank CIRMF, IRD and OMSA, for general support. We thank the people who kindly participated in our study in the different villages of the Ogooué-Ivindo. The computations presented in this article were carried out on the CALI calculator of the University of Limoges (CAlcul en LImousin), funded by the Limousin region, the European Union, the XLIM, IPAM, GEIST institutes and the University of Limoges.

## Author contributions

Conceptualization: Lokman Galal and Aurélien Mercier

Data Curation: Lokman Galal, Matthieu Fritz, Pierre Becquart, Karine Passebosc, Nicolas Plault and Aurélien Mercier

Formal Analysis: Lokman Galal and Karine Passebosc

Funding Acquisition: Lokman Galal, Aurélien Mercier and Eric M. Leroy

Investigation: Lokman Galal, Karine Passebosc and Nicolas Plault Methodology: Lokman Galal

Project Administration: Lokman Galal and Aurélien Mercier

Resources: Matthieu Fritz, Pierre Becquart, Linda Bohou Kombila, Telstar Ndong Mebaley, Larson Boundenga, Illich Manfred Mombo, Léadisaelle Hosanna Lenguiyah, Barthélémy Ngoubangoye, Nadine N’Dilimabaka, Eric M. Leroy and Gael Darren Maganga

Software: Lokman Galal

Supervision: Lokman Galal and Aurélien Mercier

Validation: Matthieu Fritz, Pierre Becquart, Linda Bohou Kombila, Telstar Ndong Mebaley, Larson Boundenga, Illich Manfred Mombo, Léadisaelle Hosanna Lenguiyah, Barthélémy Ngoubangoye, Nadine N’Dilimabaka, Eric M. Leroy and Gael Darren Maganga

Visualization: Lokman Galal

Writing – Original Draft Preparation: Lokman Galal

Writing – Review & Editing: Lokman Galal and Aurélien Mercier

